# Engineering resistance against *Tomato yellow leaf curl virus* via the CRISPR/Cas9 system in tomato

**DOI:** 10.1101/237735

**Authors:** Manal Tashkandi, Zahir Ali, Fatimah Aljedaani, Ashwag Shami, Magdy M. Mahfouz

## Abstract

CRISPR/Cas systems confer molecular immunity against phages and conjugative plasmids in prokaryotes. Recently, CRISPR/Cas9 systems have been used to confer interference against eukaryotic viruses. Here, we engineered *Nicotiana benthamiana* and tomato *(Solanum lycopersicum)* plants with the CRISPR/Cas9 system to confer immunity against the *Tomato yellow leaf curl virus* (TYLCV). Targeting the TYLCV genome with Cas9-single guide RNA at the sequences encoding the coat protein (CP) or replicase (Rep) resulted in efficient virus interference, as evidenced by low accumulation of the TYLCV DNA genome in the transgenic plants. The CRISPR/Cas9-based immunity remained active across multiple generations in the *N. benthamiana* and tomato plants. Together, our results confirmed the efficiency of the CRISPR/Cas9 system for stable engineering of TYLCV resistance in *N. benthamiana* and tomato, and opens the possibilities of engineering virus resistance against single and multiple infectious viruses in other crops.

## Introduction

Tomato (*Solanum lycopersicum*) is an important horticultural crop grown worldwide, with a global production of more than 159 million metric tons on nearly 50 million hectares, and a global trade value of 16.5 billion USD in 2010 (FAO statistics 2010–11). However, tomato is susceptible to a range of pathogens, including viruses [1]. *Tomato yellow leaf curl virus* (TYLCV) is the most economically important monopartite virus of the genus *Begomovirus* of the Geminiviridae family, and TYLCV can cause 100% crop loss in the field [2]. Tomato plants infected with TYLCV show severe symptoms of stunting with small, thick, rubbery cup-shaped leaves and yellowing along leaf margins, which leads to significant fruit loss [3]. First detected in the Middle East in 1960, TYLCV is now endemic in many parts of the world, including Africa, The Americas, Asia, and Australia [4]. TYLCV is transmitted by an insect vector, *Bemisia tabaci* (whitefly).

TYLCV is composed of twin icosahedral capsids with a circular, single-stranded DNA genome of 2.7 kb that encodes six to seven partially overlapping open reading frames (ORFs) in bidirectional organization for viral replication and spread [5]. Two of these ORFs (V1 and V2) are present in the virion sense orientation and four of them (C1–C4) are in the antisense orientation separated by a 300-nucleotide (nt) intergenic region (IR). The IR contains the key element responsible for replication and transcription of the viral genome. The virion sense ORF V1 encodes the viral coat protein (CP, 30.3 kDa), which is responsible for encapsidation of the genome and is involved in virus movement and vector recognition (15). ORF V2 (13.5 kDa) encodes the pre-coat protein, which is involved in viral spread from cell to cell (n5). The replication-associated protein (Rep, 41 kDa) is encoded by ORF C1 located in the antisense orientation of the viral genome. ORF C2 (15.6 kDa) encodes a transcriptional activator protein that is involved in the activation of transcription and ORF C3 (15.9 kDa) and ORF C4 (10.9 kDa) encode proteins involved in viral DNA accumulation [6].

Multiple approaches are used to control TYLCV and other viruses in crops, including the use of resistant cultivars, the introduction of resistance (R) genes, RNA interference (RNAi), recessive genomic mutational tactics, and pesticides to control vectors [7-11]. However, the ability of the viruses to overcome the endogenous or engineered control measures poses a challenge in the eradication of crop viruses. In agriculture, the susceptibilities of preferred varieties, the use of mixed cropping systems, susceptible plant developmental stages, and the presence of mixed viral strains, conducive environments, and transmission vectors further complicate viral disease management [3].

Clustered regularly interspaced short palindromic repeats and associated proteins (CRISPR-Cas) systems are emerging as a novel method to engineer virus resistance. *Streptococcus pyogenes* CRISPR-Cas9 is the simplest and most studied system, and the endonucleolytic complex (Cas9-guide RNA (gRNA)-tracer RNA (tracRNA) is programmable for the site-specific cleavage of dsDNA. Any target in the DNA sequence followed by NGG (the protospacer adjacent motif, PAM) can be cleaved by the Cas9-sgRNA (engineered single-guide RNA) complex. The 20 nt sequence of the sgRNA complementary to the target DNA works as a guide and the stem-loop scaffold binds and activates Cas9 for sequence-specific nuclease activity [12, 13]. The easy engineering of the CRISPR-Cas9 system allows its application as an RNA-guided DNA endonuclease for a variety of genome engineering applications across diverse eukaryotic species, including as an effective platform to generate resistance against viruses, such as DNA viruses that infect plants [14-16].

Previous reports have shown the utility of this system to confer resistance against DNA viruses in transient assays in model plant species [14]. Here, we demonstrated that the CRISPR-Cas9 system can target the TYLCV genome in tomato plants resulting in efficient virus interference. Specifically, plants expressing sgRNA targeting the CP sequence of TYLCV showed better viral interference compared to those targeting the Rep sequence of the TYLCV genome. Additionally, we showed that the CRISPR/Cas9 machinery-mediated interference was inherited through multiple generations, thereby demonstrating its utility for developing durable virus resistance. Moreover, our results demonstrated that targeting the coding regions of the virus may lead to the generation of variants capable of overcoming the CRISPR/Cas9 machinery. However, simultaneous targeting of multiple coding and non-coding virus sequences results in effective interference. Together, our results confirmed that the CRISPR/Cas system can be introduced into plants for engineering durable virus resistance against TYLCV.

## Results

### Engineering of CRISPR/Cas9-mediated immunity against TYLCV in *N. benthamiana*

Recently, we established an efficient method to target the genomic DNA of different viruses using CRISPR/Cas9 [14] and confirmed that targeting different coding and non-coding sequences of geminiviruses results in effective viral interference [17]. To test whether CRISPR-Cas9 systems can be used to confer durable virus resistance *in planta*, we engineered plants that reliably express the CRISPR/Cas9 machinery targeting TYLCV. We used *Agrobacterium-mediated* T-DNA transformation to express sgRNAs from the *U6-26s* promoter and *Cas9* under the control of the *CaMV-35S* promoter in the model plant *N. benthamiana.* The *U6-sgRNA* cassette and the human codon-optimized *Cas9* gene under the *CaMV-35S* promoter were cloned into a binary vector (Figure 1A) and transformed into *N. benthamiana* leaf discs using *Agrobacterium tumefaciens.* The primary transformants were selected on regeneration media with kanamycin as the selection marker and transferred to soil. The presence of the Cas9 endonuclease was confirmed in three individual transgenic lines by western blotting with an antiFLAG antibody (Supplementary Figure 1).

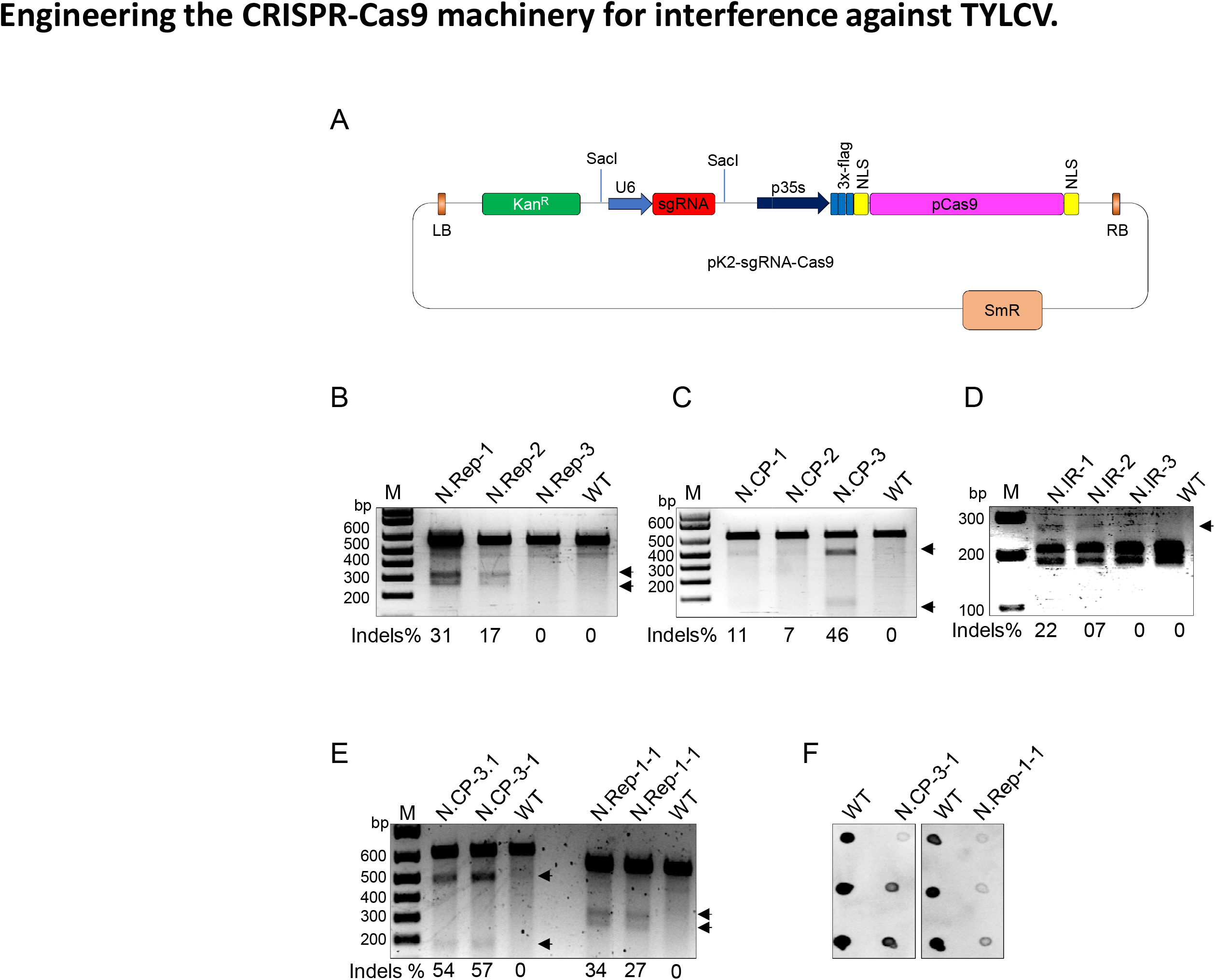
Engineering the CRISPR-Cas9 machinery for interference against TYLCV.

To confirm that lines expressing CRISPR/Cas9 were capable of targeting and cleaving the viral genome, the infectious clone of TYLCV in a binary vector was introduced into the lower leaf of the transgenic and wild-type plants through agro-infiltration. Total DNA was isolated from the systemic leaves at seven days post infiltration. The target-encompassing region was PCR amplified using TYLCV-specific primers (Supplementary Table 1) and subjected to the T7EI assay for the detection of insertions/deletions (InDels). Our results clearly demonstrated the ability of the CRISPR-Cas9 system to target TYLCV at the CP, IR, or Rep sequences in *N. benthamiana* plants (Figure 1B, C and D).

Silencing of the T-DNA inserted genes is common in plants, particularly when these genes are engineered for viral resistance [18]. Therefore, we tested the capability of the system to confer virus interference throughout multiple generations. To this end, we inoculated T_3_ progeny plants with TYLCV and conducted molecular analysis to test for virus interference and modification. T7EI (Figure 1E) and dot-blot analyses (Figure 1F) confirmed that TYLCV was effectively targeted by the CRISPR/Cas9 system in the T**3** plants, and this targeting resulted in low accumulation of the TYLCV genome in the plants, which was similar to the results of the T_1_ generation.

### A stably-engineered CRISPR/Cas9 system conferred interference against TYLCV in tomato

To engineer CRISPR-Cas9-based resistance against TYLCV in tomato, the T-DNA of the binary constructs (Figure 1A) was transformed into tomato cotyledons (cultivar Money Maker) and transgenic lines expressing CRISPR-Cas9 machinery were regenerated. Six individual lines (for each of the CP and Rep targets) were confirmed for the expression of Cas9 by anti-flag antibodies (Supplementary Figure 2) and were grown to maturity to collect T_2_ seeds. T_2_ seedlings were selected on kanamycin, acclimatized and transferred to soil, and then inoculated with TYLCV. Total DNA was isolated from leaves for various molecular analyses. PCR amplicons encompassing the target regions of the CP and Rep regions of the TYLCV genome were subjected to T7EI mutation detection analysis. Our T7EI results confirmed efficient targeting of the CP and Rep sequences in the plants expressing CRISPR-Cas9 (Figure 2A and B). Subsequently, we conducted Sanger sequencing of the PCR amplicons and validated the targeting of TYLCV by the Cas9 endonuclease in the T_2_ transgenic plants (Figure 2C and D).

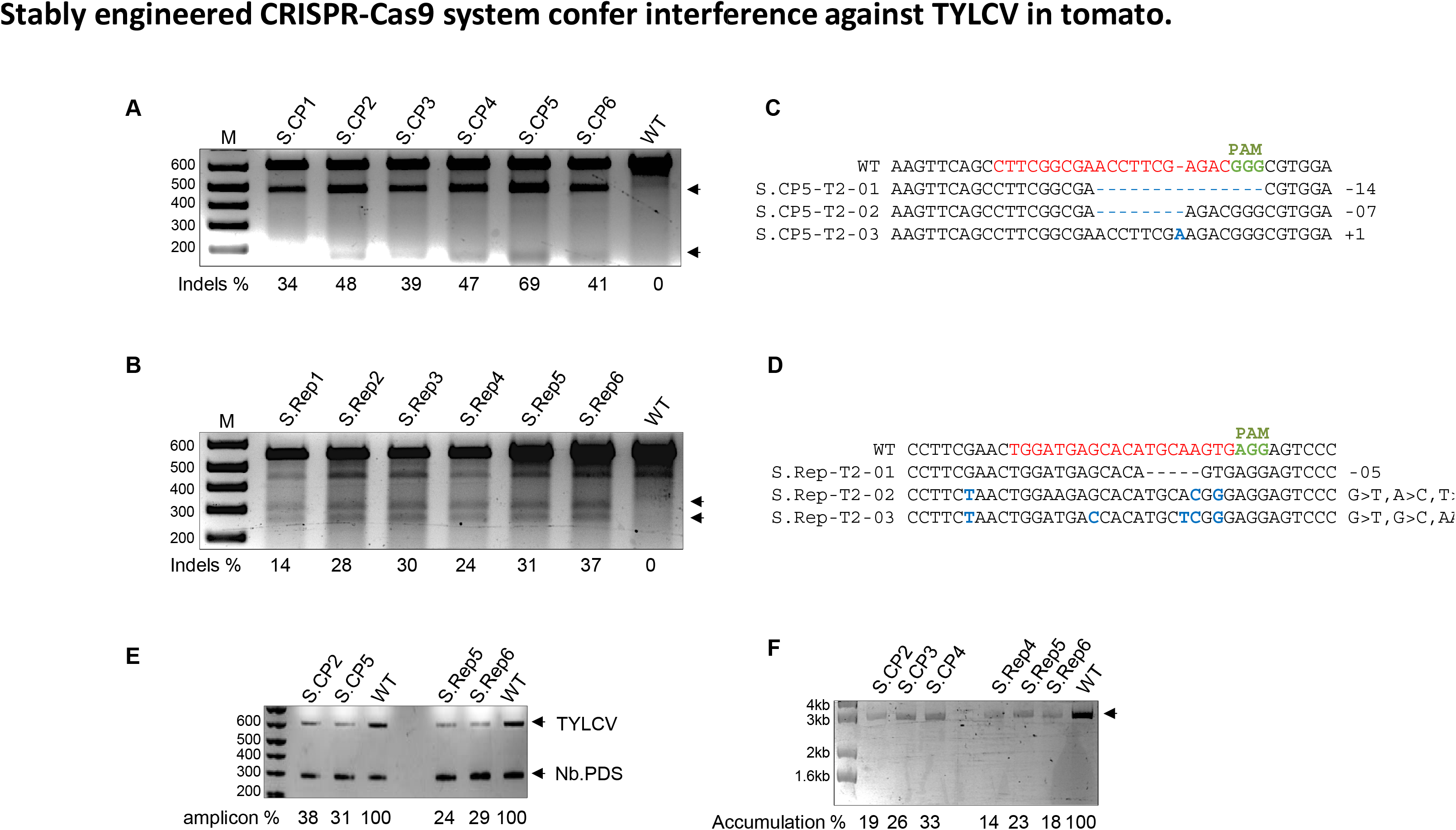
Stably engineered CRISPR-Cas9 system confer interference against TYLCV in tomato.

To confirm that targeting by CRISPR-Cas9 results in low accumulation of the TYLCV genome in tomato, extracted DNA was subjected to semi-quantitative PCR analysis. Transgenic plants accumulated a lower titer of the TYLCV genome compared to the wild-type plants (Figure 2E). TYLCV is a single-stranded DNA virus and converts to double-stranded DNA by rolling circle amplification. To test whether TYLCV targeting via the CRISPR/Cas9 machinery in transgenic plants resulted in low accumulation of the single-stranded viral DNA genome, we performed a rolling circle amplification assay (RCA). The RCA assay results showed that the transgenic plants accumulated a lower amount of viral genomic DNA compared to the wild-type plants (Figure 2F).

### CRISPR/Cas9-mediated immunity against TYLCV was stable over multiple generations in tomato

To engineer viral immunity in crops like tomato, the function of the CRISPR/Cas9 system must be inherited indefinitely. To validate the inheritance of the CRISPR/Cas9 machinery and function in virus interference, T_3_ progeny plants were inoculated with TYLCV and various molecular analyses were conducted on the extracted DNA. Our T7EI results confirmed efficient targeting of the CP and Rep sequences in the T_3_ plants expressing the CRISPR/Cas9 machinery, indicating its ability to confer virus resistance in the progeny plants (Figure 3A and B). Further, our Sanger sequencing data of the PCR amplicons flanking the target sequence in the virus genome confirmed effective targeting of TYLCV by the Cas9 endonuclease (Figure 3C and D).

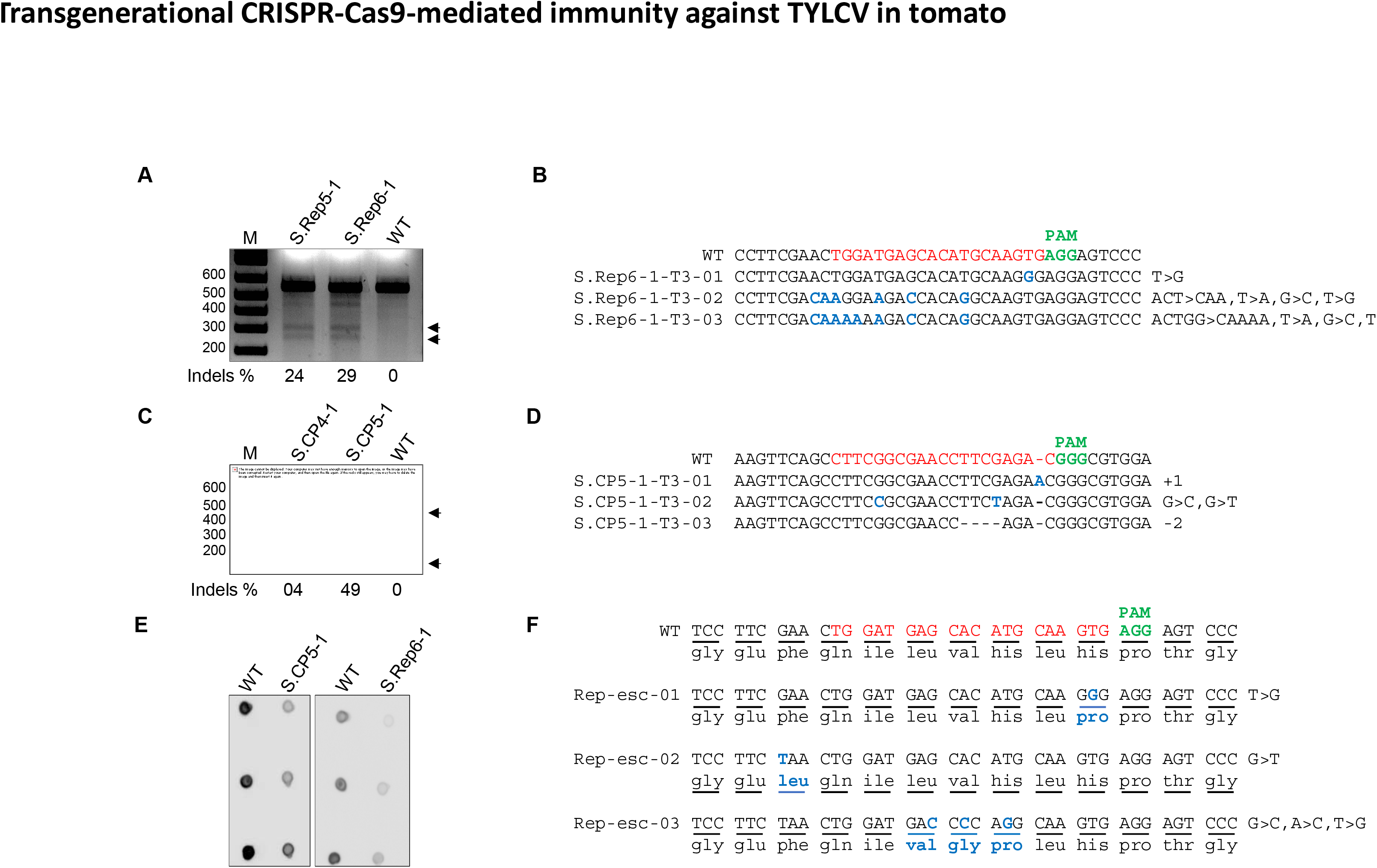
Transgenerational CRISPR-Cas9-mediated immunity against TYLCV in tomato

To confirm that CRISPR-Cas9-mediated immunity results in a lower titer accumulation of TYLCV, we conducted semi-quantitative PCR and RCA analysis on T**3** progeny plants inoculated with TYLCV. Both the semi-quantitative PCR (Supplementary Figure 3) and RCA data (Supplementary Figure 4) clearly demonstrated low accumulation of the TYLCV genome in the T_3_ homozygous plants expressing the CRISPR-Cas9 system. We used dot-blot analysis to determine the virus titer, which indicated a lower accumulation of TYLCV genomic DNA in the T_3_ homozygous plants than in the wild-type control plants (Figure 3E). These data indicated that the CRISPR-Cas9 system engineered in plants to target the TYLCV provides an effective tool to control virus infection.

### TYLCV evasion from the CRISPR/Cas9 machinery

Viruses could use the repair machinery to generate variants capable of escaping the CRISPR/Cas9 machinery. For example, TYLCV genome can be targeted by CRISPR/Cas9 machinery for cleavage and can be repaired by cellular NHEJ repair pathway in plant cell nucleus. Such imprecise repair could allow TYLCV genome to evolve and change the sequences corresponding to the spacer or protospacer-associated motif, thereby evading the CRISPR/Cas9 system and replicating and spreading systemically. It is worth noting that these virus escapees would be un-targetable and able to freely replicate and move across the plants. To understand and limit the frequency of these escapees are key to develop durable virus resistance.

To test the ability of TYLCV to overcome the CRISPR-Cas9 system, we collected sap from the TYLCV-infected *N. benthamiana* transgenic plants expressing CRISPR/Cas9 targeting the TYLCV Rep sequence and used this sap to mechanically inoculate wild-type *N. benthamiana* plants. DNA was collected 7 days after inoculation (dai) and subjected to rigorous molecular analysis. Sanger sequencing confirmed the Rep sequence variants of the TYLCV genomes and analysis of the modified nucleotide sequence to their corresponding amino acids demonstrated that, almost all of evolved variants have functional and contiguous ORFs but with varied amino acids at targeted sequence (Figure 3F) thereby allowing replication and systemic movement.

## Discussion

CRISPR/Cas9 is an efficient, adaptive anti-viral system of bacteria and archaea that was recently tailored for targeted modifications of eukaryotic genomes and to provide interference against viruses, including DNA viruses that infect plants [17, 19, 20]. The CRISPR/Cas9 system can be introduced into plants for interference against DNA viruses and can also be used to mutate a susceptible gene in the plant genome to confer resistance [14]. TYLCV is one of the most serious viruses infecting tomato and other important crops, but conventional methods to control TYLCV are expensive and mostly ineffective [3]. Here, we assessed the utility of the CRISPR/Cas9 system to confer resistance against TYLCV in *N. benthamiana* and tomato, and tested the durability of virus resistance over multiple generations. Most importantly, we also assessed whether the CRISPR/Cas9 machinery would trigger mutations in the viral genome capable of evading the CRISPR/Cas9 system.

We stably engineered the CRISPR/Cas9 machinery in *N. benthamiana* and tomato to target TYLCV genomic sequences. CRISPR/Cas9 targeted the CP sequence of TYLCV very efficiently and provided robust interference in all tomato plants from the T_2_ to the homozygous T_3_ generation.

The CRISPR/Cas9 system also targeted the Rep region of the viral genome, though with less efficiency compared to the CP sequence. One possible explanation for this difference in efficiency is that the sgRNA directing the Cas9 endonuclease is more efficient in binding to the CP target sequence than to the Rep sequence. The presence of binding proteins or competition with the viral replication machinery for binding to the *Rep* ORF could hinder the binding of the CRISPR-Cas9 complex to the spacer sequence in the *Rep* region. The latter possibility is consistent with recent data showing the robust targeting and cleavage of the CP sequence of another geminivirus, *Cotton leaf curl kokran virus* (CLCuKV), by the CRISPR/Cas9 machinery [17]. Interestingly, all possible Indels were detected, but we observed in the targeting of TYLCV in permanent lines of *N. benthamiana* and tomato expressing the CRISPR/Cas9 system, a predominant single nucleotide change at the Cas9 targeted sequence.

Previous observations have indicated that silencing of the CRISPR-Cas9 system can occur in successive generations [18, 21]. Therefore, after demonstrating the efficiency of stably-expressed CRISPR/Cas9 machinery in conferring virus interference, we tested whether this activity persists over multiple generations. Indeed, our results demonstrated that the expression of the CRISPR/Cas9 machinery and viral resistance was inherited over multiple generations.

Viruses can often overcome the plant’s genetic immunity, including the engineered CRISPR/Cas9 system [17]. It is crucial to understand the limits and frequency of this naturally-acquired resistance in order to develop durable virus resistance in plants [22]. Additionally, targeting the viral genome by the CRISPR/Cas9 system leads to the formation of double-strand breaks, which are repaired by error-prone non-homologous end joining or more precise homology-directed repair mechanisms. Thus, this repair machinery could generate viral variants capable of overcoming the CRISPR/Cas9 machinery, as imprecise repairs of the viral genome could result in mutations in the sequences corresponding to key sequences essential for the Cas9 activity including spacer and PAM sequences. Because viruses have high rates of evolution and the CRISPR/Cas9 system does not tolerate mismatches in the seed sequences (9–12 nt) of the spacer near the PAM sequence [23], any mutation of these sequences would limit or eliminate the ability of the CRISPR/Cas9 system to target the virus, and therefore lead to virus variants that can overcome the resistance. To test this, we assessed the ability of the TYLCV genome to overcome the CRISPR/Cas9-mediated immunity. Similar to our previous results for CP [14] Sanger sequencing data revealed TYLCV genomes with sequence modifications at the Rep sequence, but maintaining a functional and contiguous ORFs, thereby capable of replication and systemic movement.

Intergenic region nona nucleotides (IR) important for virus replication can limit the frequency of these viral escapees [14]. However, multiple attempts to regenerate tomato lines expressing Cas9 and sgRNA targeting the IR failed, whereas we were able to successfully regenerate tomato lines with Cas9 and sgRNA targeting the CP and Rep regions. Alternative approaches including the simultaneous targeting of the virus at multiple sequences could boost resistance and limit the chances of virus variants capable of replication [24]. Protein binding to the TYLCV genomic DNA can hinder the binding of the viral replication machinery [25]. For example, a dCas9 approach to allow binding of multiple sgRNA-dCas9 complexes to the viral genome may interfere with the replication machinery leading to effective virus interference. Recently, several new mechanisms of viral interference were discovered, including CRISPR/Cas13a/b systems which target the virus RNA genome or the RNA intermediate of a DNA virus [26, 27]. Developments in CRISPR/Cas systems are opening new possibilities to engineer durable virus resistance in crops and improve crop yield and productivity.

## Material and methods

### Vector construction

To generate tomato and *N. benthamiana* lines expressing Cas9 and sgRNA, the U6-sgRNA and *Cas9* were assembled into the binary pK2GW7 vector. First, the 20-nt spacer sequences of the IR, CP, or Rep sequences (singly or tRNA-sgRNA) of the TYLCV genome were cloned into a backbone (U6-BbsI-gRNA scaffold) vector as primer dimer by Golden Gate enzyme *Bbs*I. Next, the whole U6-sgRNA cassette was amplified with a forward and reverse primer set both containing a SacI digestion site. By restriction ligation, the SacI-U6-sgRNA-SacI fragment was moved to pK2GW7. Next, the 3XFlag–NLS-Cas9-NLS cassette was cloned into the pENTR/D-TOPO plasmid (Life Technologies) and moved to the U6-sgRNA-pK2GW7 destination vector by Gateway LR clonase in front of the CaMV-35S promoter to make the U6-sgRNA-pK2-Cas9 T-DNA vector (Life Technologies) [28].

### Generation of tomato and *N. benthamiana* lines expressing *Cas9* and sgRNA

The U6-sgRNA-pK2.Cas9 T-DNA binary vector was introduced into *A. tumefaciens* strain GV3101 by electroporation and transgenic *N. benthamiana* plants were regenerated following a previously described method [29]. To regenerate tomato lines expressing sgRNA and *Cas9*, the T-DNA construct was transformed into tomato (cultivar Money Maker) cotyledons and plants were regenerated using a previously described method [30]. Small transgenic plantlets were moved to soil for acclimation. The expression of Cas9 in T_1_, T_2_, and T_3_ plants was confirmed by western blot using an anti-Flag antibody [29].

### T7EI mutation detection assay

To determine the Cas9 endonuclease activity, the non-homologous end joining-based double-strand break repair rate was evaluated by the T7EI assay. Total genomic DNA was isolated from samples collected after 7 days (for *N. benthamiana*) and 15 days (for tomato) and used as a template to amplify the respective spacer flanking fragments with individual primer sets using Phusion polymerase. Next, 200 ng of the PCR product was subjected to the T7EI assay [29]. ImageJ software (http://rsb.info.nih.gov/ij) was used to estimate the mutation rates. The PCR products were cloned into the pJET2.1 cloning vector for Sanger sequencing.

### TYLCV infections

*Agrobacterium* containing the infectious T-DNA clone of TYLCV (TYLCV2.3) was infiltrated into young lower leaves of *N. benthamaina* or cotyledons of tomato plants expressing the CRISPR-Cas9 machinery. For sap inoculation, the leaf tissue was mashed in plastic bags with 10 ml of Tris-phosphate buffer (pH 7.5) and rub-inoculated into young leaves of *N. benthamiana* or cotyledons of tomato using corborandom (mesh size 200–400 nm).

### Determination of viral variants

The sap inoculation approach was used to screen for potential viral variants. Sap was collected form the TYLCV-infected CRISPR/Cas9-expressing plants and was mechanically applied to wild-type *N. benthamiana* plants. At 7 dai, samples were collected and DNA was extracted and the respective fragments were PCR amplified for cloning into pJET2.1 for Sanger sequencing.

### Search for potential off-target of IR spacer sequence

These spacer binding sites were used to find the exact match or single mismatches in the genome of tomato (*S. lycopersicum*) http://solgenomics.net/. The putative sites were subjected to further annotation, where sequences were split into two groups of exact match and off-target binding (1–7 mismatches) in the seed sequences.

### Rolling circle amplification

Genomic DNA (50 ng) isolated from CRISPR-Cas9-expressing plants infected with TYLCV was subjected to a rolling circle amplification (RCA) assay using the RCA amplicon kit following the manufacturer’s protocol (GE Healthcare). The DNA samples were resolved on a 1% agarose gel.

## ACKNOWLEDGMENTS

We thank the members of the Laboratory for Genome Engineering for their continuous constructive discussions and comments on the manuscript. This publication is based upon work supported by the King Abdullah University of Science and Technology (KAUST) Office of Sponsored Research (OSR) under Award No. OSR-2015-CRG4-2647.

## COMPETING FINANCIAL INTERESTS

The authors declare no competing financial interests.

## Author Contributions

MM conceived research. MT, ZA, FA, AS performed research, MT, ZA, FA, AS and MM analyzed data, ZA and MM wrote the paper.

